# Accurate Site-specific Folding via Conditional Diffusion Based on Alphafold3

**DOI:** 10.1101/2025.07.06.663385

**Authors:** Haocheng Tang, Junmei Wang

## Abstract

Accurate structure prediction of biomolecular complexes is crucial for understanding biological processes and enabling drug discovery. While AlphaFold3 represents a significant advancement, enhancing its accuracy for specific binding sites remains a challenge. We present SiteAF3, a novel method for accurate site-specific folding via conditional diffusion, built upon the AlphaFold3 framework. SiteAF3 refines the diffusion process by fixing the receptor structure and optionally incorporating binding pocket and hotspot residue information. Comprehensive evaluations on protein-small molecule, protein-peptide, and protein-nucleic acid datasets demonstrate that SiteAF3 consistently outperforms AlphaFold3, achieving higher accuracy in complex structure prediction especially for orphan proteins and allosteric ligands, with reduced computational cost. SiteAF3 offers a user-friendly plug-in compatible with AlphaFold3, providing a valuable tool for more accurate modeling of biomolecular interactions.

## 1 Introduction

Accurate structure prediction of biomolecular complexes is fundamental to understanding biological processes and essential in rational drug discovery and development. The revolutionary advancements in artificial intelligence, particularly with the introduction of AlphaFold2 (AF2) [1], marked a significant leap forward in predicting the three-dimensional structures of individual proteins with unprecedented accuracy. Building upon this success, Alphafold3 (AF3) represents the next major milestone [2], expanding its capabilities to encompass a far broader range of biomolecules and their interactions.

Despite its remarkable progress, AF3 still presents certain limitations. One primary challenge is that its performance heavily depends on multiple sequence alignment (MSA), which can be problematic for orphan proteins with limited sequence homology or when the generated MSA contains errors, potentially leading to inaccurate predictions [2–5]. Furthermore, the co-folding model has difficulty accurately predicting the binding of allosteric ligands [6]. Additionally, significant improvements can still be made in the accuracy of both predicted interfaces and overall complex structures. For computational cost, the runtime for conformational sampling is still much slower compared to traditional docking methods, and running AF3 on large biomolecular systems may encounter difficulty related to GPU memory [7]. Moreover, the pocket-guided prediction functionality in AF3 has not been made publicly available, limiting its accessibility and customizability for specific applications.

To address these limitations and further enhance the accuracy of site-specific complex structure prediction, we introduce SiteAF3, an innovative tool for accurate site-specific folding via conditional diffusion based on AF3. SiteAF3 builds upon the foundation of AF3 by incorporating two key modifications. Firstly, we fine-tune the original diffusion module of AF3, implementing a conditional diffusion process where the receptor structure is fixed. This allows for a more focused exploration of the conformational space of the ligand in the context of a pre-defined receptor. Secondly, SiteAF3 offers the option to explicitly guide the prediction process by incorporating information about the binding pocket and/or hotspot residues through the MSA module. By providing SiteAF3 with the experimentally determined or high-confidence predicted structure of the receptor and pocket information, our method generates accurate structures of the ligand-receptor complex. SiteAF3 offers several notable advantages and features:

- **High accuracy** Our results demonstrate that it achieves higher accuracy in predicting biomolecular complex structures compared to the original AF3, especially for MSA-helpless orphan structures and allosteric binding ligands prediction.
- **High compatibility** SiteAF3 is designed for seamless integration and ease of installation as a lightweight plug-in for AF3. SiteAF3 maintains the broad compatibility of AF3, supporting the prediction of complexes involving proteins, small molecules, nucleic acids, and ions, and it can handle multiple chains within the complex.
- **Low computational cost** SiteAF3 exhibits lower GPU memory consumption for large complex systems, particularly when the token number of fixed chains in receptors increases. Besides, it offers significantly faster prediction speeds compared to the standard AF3 pipeline when not using genetic searching, allowing for rapid high-throughput virtual screening.

In this work, we present comprehensive benchmarking results and case studies demonstrating the performance of SiteAF3. We show that our method consistently outperforms AF3 across diverse datasets of small molecules, peptides, double-stranded DNA (dsDNA), and RNA molecules interacting with proteins. These results highlight the efficacy of our conditional diffusion approach and the benefits of incorporating site-specific information. In summary, SiteAF3 represents a significant advancement in accurate site-specific folding of biomolecular complexes, offering improved accuracy, efficiency, and versatility compared to AF3 and opening new avenues for traditional docking in co-folding era.

## 2 Results

### 2.1 Network architecture of SiteAF3

The overall architecture of SiteAF3 (Fig. 1a) mirrors that of AlphaFold3 (AF3), inheriting AF3’s inference workflow, which comprises four primary components: the input preparation, the representation learning, the structure prediction, and the confidence assessment. Our main divergences from AF3 lie in two key aspects: firstly, the structure prediction module employs a novel conditional diffusion model, and secondly, the representation learning incorporates additional binding pocket and hotspot residue information via the MSA module. These modifications were motivated by the objective of enhancing the accuracy of complex structure prediction especially when the receptor structure prediction is poor as well as the need to addressing the specific binding site of a receptor.

**Fig. 1:**
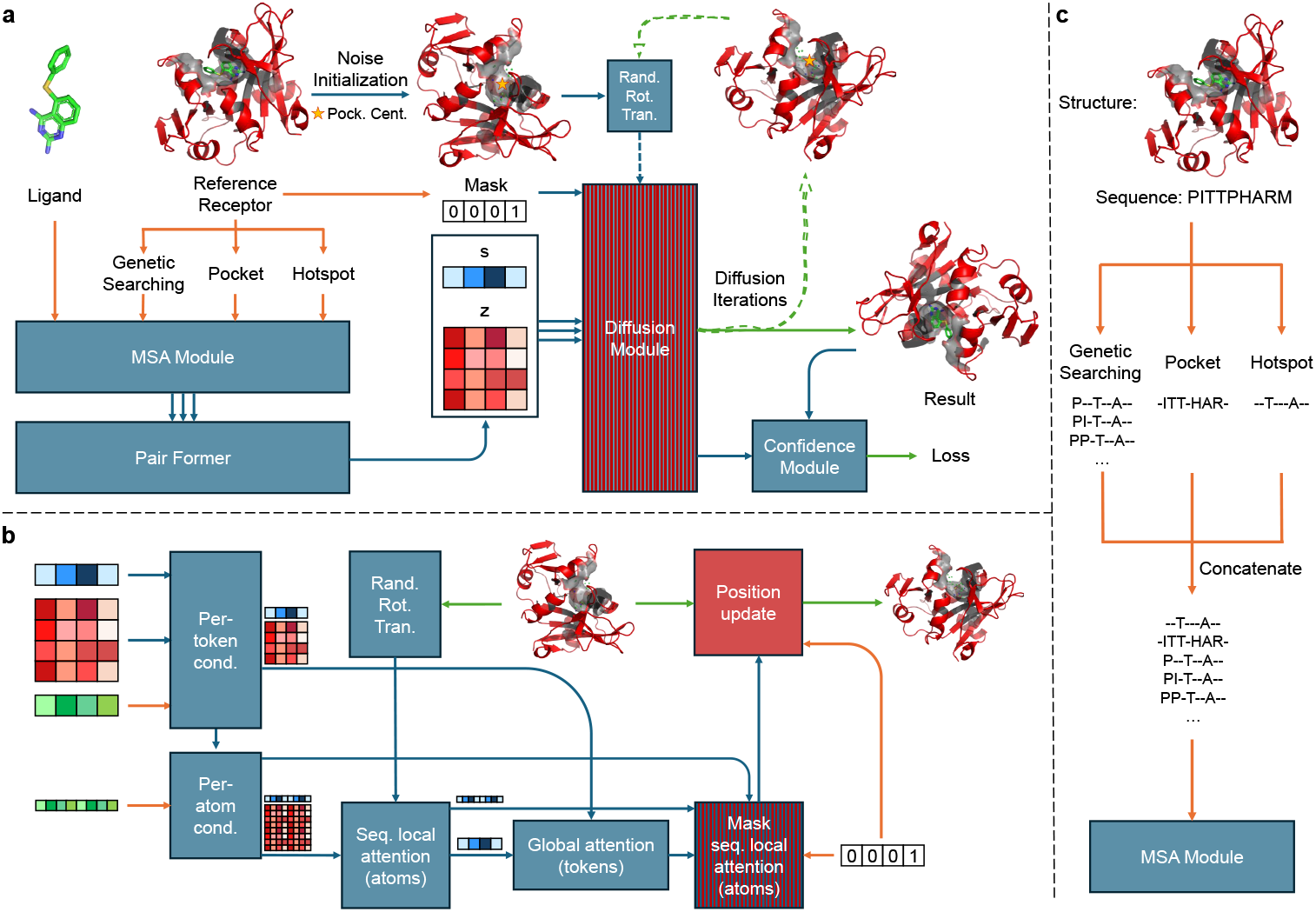
Architectural details of SiteAF3. The rectangles represent processing modules and the arrows show the data flow. Rectangles: red, new modules; blue and red stripes, fine-tuned; blue, same as in Alphafold3. Arrows: orange, input data; blue, abstract network activations; green, output data. The green dots around pockets represent physical atom coordinates. **a**, General architecture for inference. The yellow star stands for the pocket center. **b**, Conditional diffusion module with coarse arrays depicting per-token representations (green, inputs; blue, pair; red, single; 01, mask) and fine arrays depicting per-atom representations. **c**, Pocket and hotspot embedding. Abbreviations: Pock. Cent.-pocket center; Cond.-conditioning; Rand. Rot. Trans.-random rotation and translation; Seq.-sequence.

Consistent with AF3, the diffusion module in SiteAF3 (Fig. 1b and Algorithm 1) operates directly on raw atom represented as point clouds, alongside a coarse abstract token representation, eschewing rotational frames or any explicit equivariant processing. However, the initialization of noise differs from the original diffusion module. In SiteAF3, the atomic coordinates of the ligand are initialized with noise based on a Gaussian distribution centered around the pocket center with a specified radius, while the relative atomic coordinates of the receptor are directly fixed. To ensure diversity in the initial states, the pocket center is also subjected to a slight perturbation within a 2 Å-radius sphere. Furthermore, to guarantee the robustness of the output structures, the initialized point clouds undergo random translation and rotation [8]. The second different point is situated within the second sequence local attention block, where SiteAF3 introduces a mask to update only the coordinates of the ligand. Notably, the inclusion of this mask also contributes to reduced GPU memory consumption during the diffusion process, thereby expanding the applicability of the method to larger systems (Extended Data Table 2). The third notable difference is the incorporation of a “Position update” module, designed to seamlessly combine the fixed receptor structure with the updated ligand structure to form the complete complex.

In our initial explorations with the base SiteAF3 model, we observed that despite the noise initialization of ligand atomic coordinates was around the pocket, the predicted ligand binding occurred at a site distant from the intended pocket occasionally. To address this, we first used genetic searching tools in AF3 to get templates for MSA module to evaluate how much binding pocket information can be learned. We then experimented with directly embedding information about designated hotspot and pocket residues via the MSA module. These strategies led to a substantial improvement in the accuracy of the model’s predictions, effectively guiding the ligand towards the desired binding site. It is important to note that our approach differs from the pocket-guided folding within AF3 which is not open-source, and consequently, the model performance of the two models varies.

### 2.2 Results of protein and small molecule complexes

Accurate protein-small molecule structure prediction holds paramount importance in the realm of drug design. While AF3 has already demonstrated superior performance compared to traditional docking tools and other deep learning methodologies in this critical area, our method, SiteAF3, propels the accuracy of protein-small molecule structure prediction to an even higher level. To rigorously evaluate the performance of SiteAF3 and compare it against AF3, we selected two benchmark datasets: Fold-Bench protein-ligand [9] (Fig. 2 and Extended Data Table 1) and PoseBustersV2 [10] (Extended Data Fig. 2 and Extended Data Table 1). The data within the FoldBench dataset was not included in the training set of AF3, whereas the whole PoseBustersV2 dataset was used to train AF3. Our evaluation encompassed four distinct configuration modes of SiteAF3: Mode 1 (AF3 MSA+po+hot) utilizes genetic searching (inheriting from AF3 MSA), pocket information and hotspot residue information; Mode 2 (AF3 MSA) solely relies on genetic searching (AF3 MSA); Mode 3 (po+hot) employs only pocket and hotspot residue information; and Mode 4 (baseline) does not apply additional information in the representation learning phase.

**Table 1:**
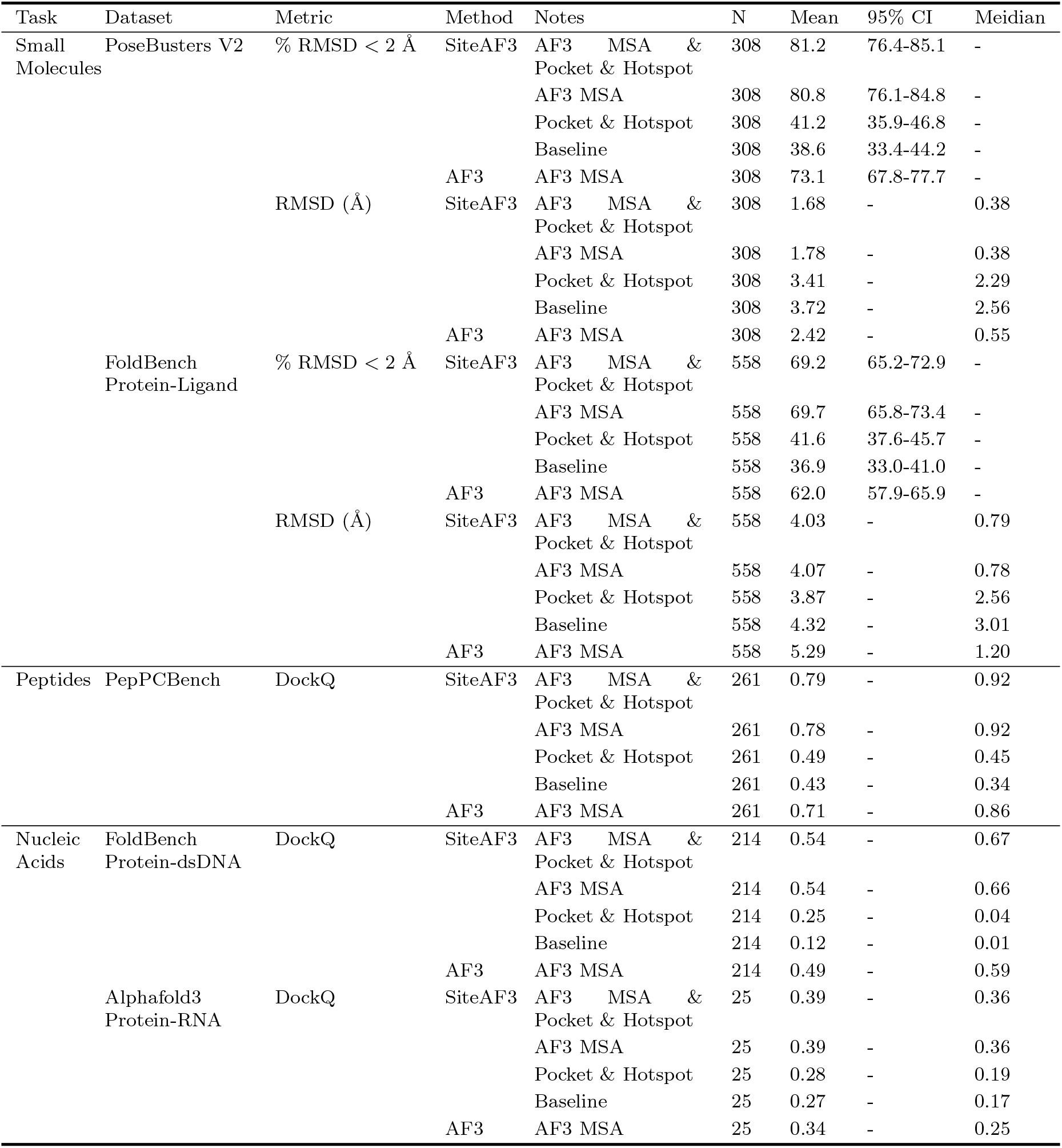
Prediction accuracy across biomolecular complexes.

**Fig. 2:**
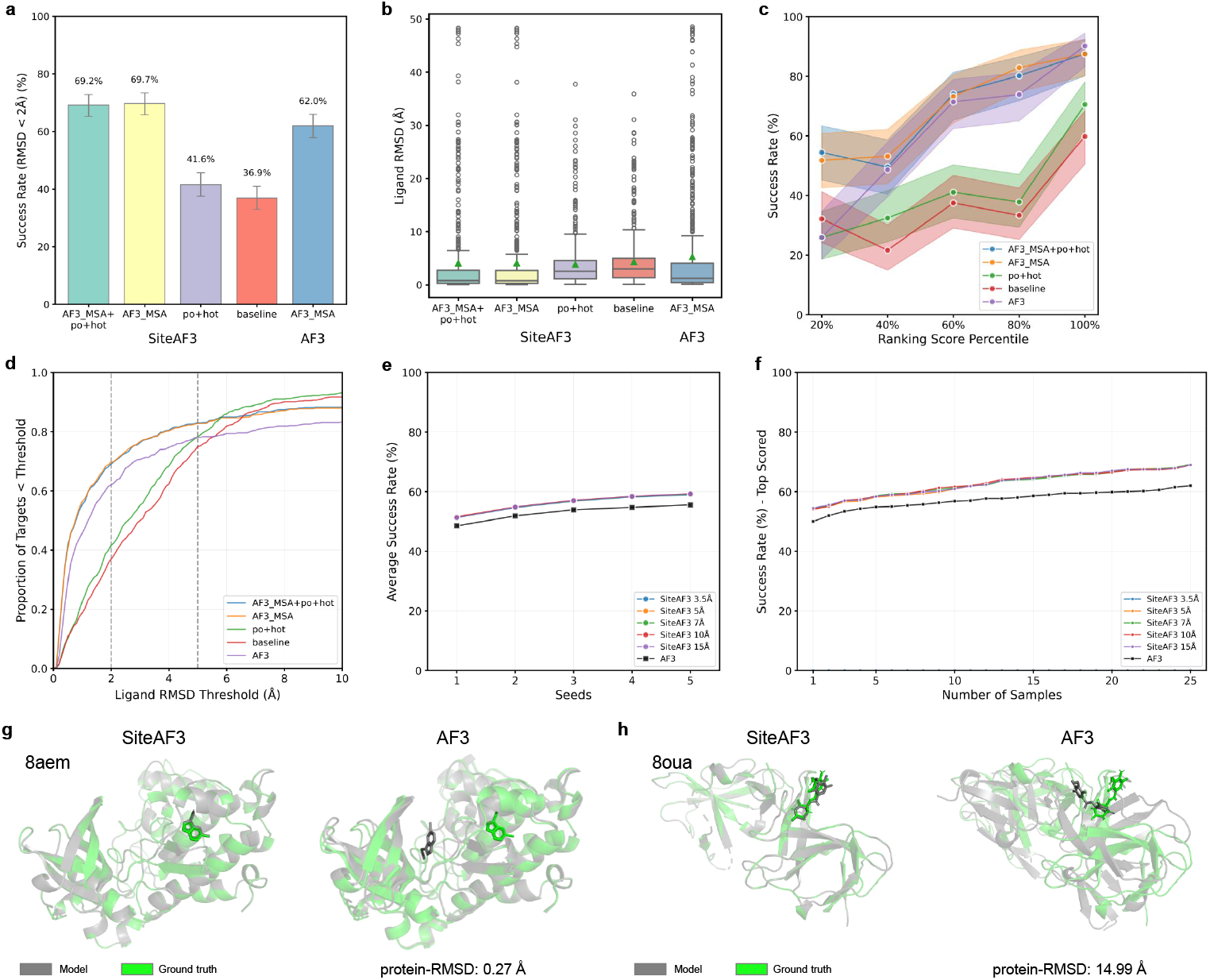
Results of protein and small molecule complexes on FoldBench protein-ligand datasets. Each sample is assigned 5 random numbers, and each random number has 5 samples. The statistic analysis adopted the best results among these 25 structures. **a**, Success rate across different subsets, where the success rate is defined as the ratio of cases with the LRMSD being less than 2 Å. **b**, Comparison of LRMSD between SiteAF3 and AF3. **c**, The relationship between success rate and confidence-related scores of SiteAF3 and AF3. The scores were segmented by number quantiles, with each bin evenly dividing our test samples. **d**, Cumulative density of LRMSD, varying from 0 to 10 Å. **e**, LRMSD average success rate by number of samples in each seed from the worst to the best seeds. **f**, LRMSD average success rates of the top-ranked samples. **g**, Case study: Allosteric compound LVF bound to CK2alpha (PDB 8aem). SiteAF3 used AF3 genetic searching, pocket and hotspot information. SiteAF3 identified the correct binding site, while AF3 successfully folded the protein but found the wrong binding pocket. **h**, Case study: cereblon isoform 4 in complex with 11F (PDB 8oua). SiteAF3 used AF3 genetic searching, pocket and hotspot information. SiteAF3 succeeded in docking while AF3 failed to generate right structure of proteins.

On the FoldBench dataset, we successfully reproduced the reported success rate of AF3 at 62.0 % (Fig. 2a), falling within the 95% confidence interval of the original paper’s reported 64.9% [9]. Notably, the best-performing SiteAF3 model achieved an accuracy of 69.7%, significantly surpassing that of AF3. Moreover, compared to the baseline model, the incorporation of pocket and hotspot information alone, without relying on AF3’s genetic searching component, also led to a substantial improvement in prediction accuracy. Examining ligand root-mean-square deviation (LRMSD) values further revealed that the best SiteAF3 model exhibited a reduction in the median by 30.9% and in the mean by 30.6% compared to AF3 (Fig. 2b and Extended Data Table 1). Fig. 2c compares the docking success rate against the ranking score. The top two performing SiteAF3 models significantly enhanced the prediction accuracy for the bottom 20% of structures with lower ranking scores, while achieving comparable performance to AF3 for structures with higher ranking scores. This indicates that SiteAF3 effectively fulfills its objective of providing more accurate predictions for structures with lower confidence scores. An intriguing result is presented in Fig. 2d, where SiteAF3, when utilizing AF3 MSA, consistently demonstrates higher cumulative prediction accuracy than the standard AF3 model within the 0-10 Å range, which aligns with our expectations. However, when not employing AF3 MSA, SiteAF3 initially underperforms compared to AF3. Interestingly, beyond a threshold of 5 Å, the proportion of correctly predicted targets for the po+hot and baseline configurations gradually exceeds that of AF3. This observation suggests that the MSA templates leveraged by AF3 might occasionally introduce misleading information regarding the binding site for unseen protein-ligand pairs. This trend was not observed in the PoseBustersV2 dataset (Extended Data Fig. 2d), where all data were included in AF3’s training set. This discrepancy suggests a potential overfitting of the genetic searching-derived data during the representation learning phase for seen datasets.

To further evaluate the practical applicability of our method, we assessed SiteAF3’s performance under scenarios mimicking real-world docking experiments where the optimal pose is often unknown. As illustrated in Fig. 2e, SiteAF3 consistently outperformed AF3 when considering the average success rate of five sampled structure in each seed across five seeds from the worst to the best. Similarly, SiteAF3 demonstrated superior performance when selecting the top-ranked structures from a total of 25 samples (Fig. 2f), highlighting its robustness even without prior knowledge of the best prediction. Furthermore, we investigated the influence of the noise initialization radius on SiteAF3’s performance on the FoldBench dataset, as shown in Fig. 2e, f and Extended Data Fig. 1a, b. Within a radius range of 3.5 to 15.0 Å, SiteAF3 exhibited higher accuracy than AF3, with no statistically significant differences observed among these radii. Extended Data Fig. 2c suggested that 5.0 Å might yield the best predictive results. However, during the inference process with radii of 5.0 Å and the smaller 3.5 Å, we encountered instances of failed structure generation due to indistinguishable atomic overlaps. Consequently, a radius of 7.0 Å was selected as the operational parameter for our model to ensure stable and reliable predictions. Finally, we note that increasing the number of seeds used for sampling can further enhance the predictive capability of SiteAF3.

For the PoseBustersV2 dataset, we did not reproduce the performance of the AF3-2019cutoff reported in the original AF3 publication, potentially due to differences in pre-processing or sampling methodologies. Nevertheless, we still observed a consistent trend where our SiteAF3 models significantly outperformed AF3 (Extended Data Fig. 2a, 2b and Extended Data Table 1). Extended Data Fig. 2c presents a different picture compared to Fig. 2c, with AF3 exhibiting a notably lower success rate across the entire range of ranking scores compared to SiteAF3. Furthermore, the ranking score demonstrated a more pronounced linear correlation with the docking success rate for SiteAF3. In contrast to Fig. 2c, the ranking score for AF3 appears to exhibit overfitting on the unseen FoldBench dataset.

Fig. 2g and Fig. 2h provide two illustrative case studies where SiteAF3 achieves higher accuracy than AF3. In the example of 8aem [11], which contains both the ATP orthosteric site and the LVF allosteric site, AF3, despite successfully generated the receptor protein structure, incorrectly placed the LVF molecule in the orthosteric site. In contrast, SiteAF3 accurately predicted the binding of LVF to the allosteric site (Fig. 2g). For another protein complex with PDB code 8oua [12], the receptor protein structure generation itself failed for AF3, consequently hindering the generation of a correct binding pose (Fig. 2h).

### 2.3 Results on protein and peptides complexes

Peptides are key mediators in up to 40% of protein-protein interactions (PPIs) [13]. Accurate modeling of protein-peptide complexes is therefore crucial for understanding fundamental biological processes and for advancing the field of peptide-based drug design. AlphaFold3 (AF3) has already shown a notable improvement in prediction accuracy for protein-peptide complexes compared to AlphaFold2-Multimer [14]. Our SiteAF3 method further enhances this prediction accuracy. We selected the Full datasets from PepPCBench as our primary benchmark for evaluation (Fig. 3 and Extended Data Table 1) [15]. Our testing included the same four configuration modes of SiteAF3 as described above.

**Fig. 3:**
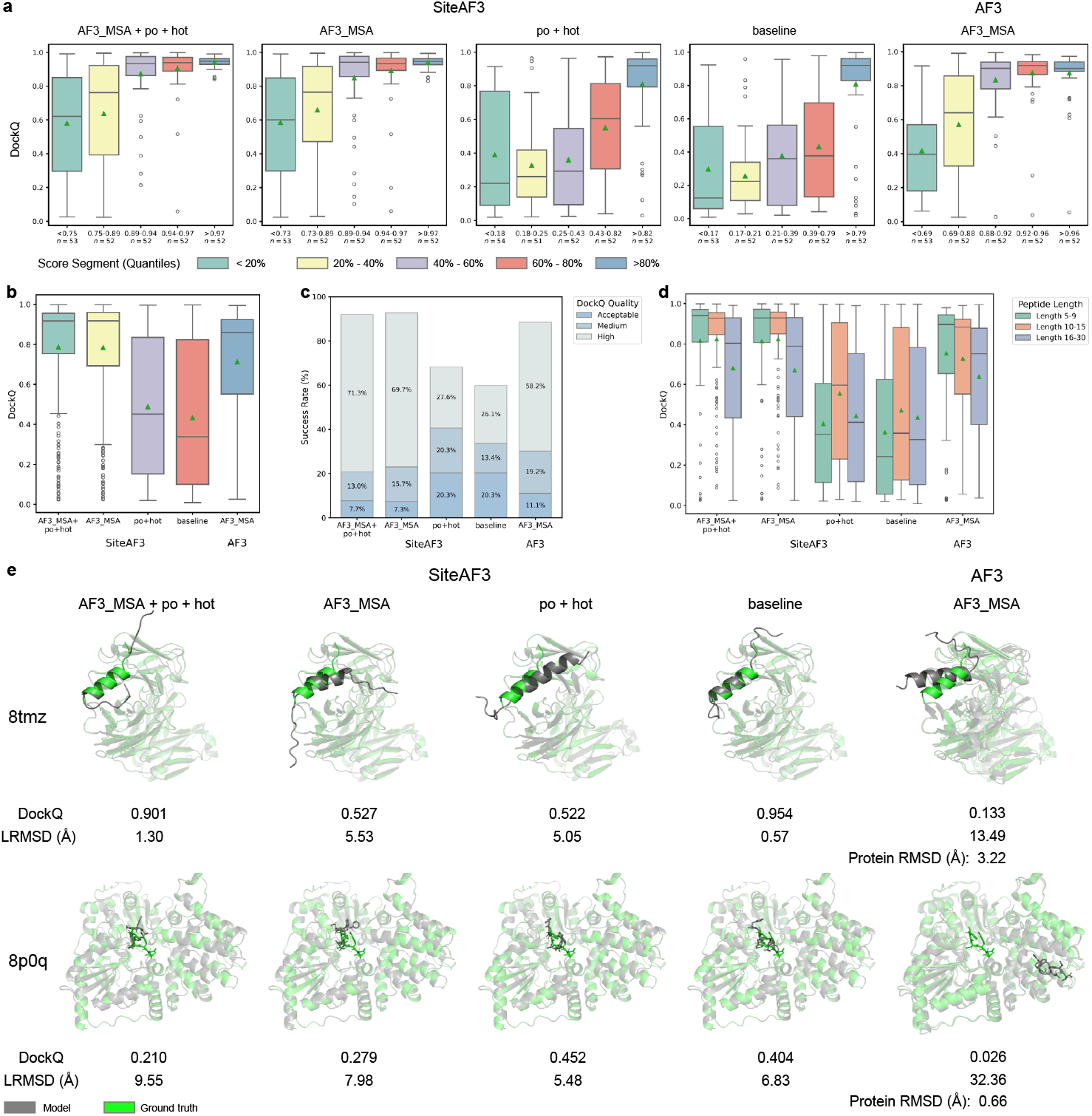
Results of protein and peptides complexes on PepPCBench datasets. Each sample is assigned 3 random numbers, and each random number has 5 samples. The statistic analysis adopted the best results among these 15 structures. **a**, The relationship between DockQ and confidence-related scores. The scores were segmented by number quantiles, with each bin evenly dividing our test samples. **b**, Comparison of DockQ value between SiteAF3 and AF3. **c**, Docking performance based on thresholds. **d**, Comparison of DockQ value across 3 different length classifications between SiteAF3 and AF3. **e**, Case studies. Up: MERS-CoV spike stem helix peptide in complex with antibody CHM-27 (PDB 8tmz). SiteAF3 succeeded in docking, while AF3 generated wrong structure of antibodies. Down: AaNGT complexed to a peptide (PDB 8p0q). SiteAF3 generated the right conformation, while AF3 successfully folded the protein but found the wrong binding site.

Our test results differ slightly from those reported in the original PepPCBench publication, which stems from their use of only the sequence corresponding to the experimentally determined structure, whereas our evaluation utilized the full-length fasta sequences available on the PDB website. Fig. 3a demonstrates that both SiteAF3 modes incorporating AF3 MSA significantly outperform AF3, particularly within the bottom 40% of structures ranked by confidence score, where SiteAF3 shows a clear lead in DockQ scores [16]. The SiteAF3 model utilizing only pocket and hotspot information also exhibits higher accuracy compared to the baseline model. Fig. 3b and Extended Data Table 1 reveal that the best-performing SiteAF3 model achieves a mean DockQ score 11.2% higher and a median DockQ score 7.0% higher than AF3. We also evaluated other key metrics such as the fraction of native contacts (fnat) [17], F1 score, interface RMSD (iRMSD), and Ligand RMSD (LRMSD) [18], as detailed in Extended Data Figure 3. These results collectively indicate that SiteAF3 is better at capturing the interactions at the protein-peptide interface and at predicting the bound conformations of highly flexible peptides. Fig. 3c shows that both SiteAF3 modes incorporating AF3 MSA exhibit a higher overall success rate compared to AF3 and, importantly, yield a significantly larger proportion of high DockQ quality predictions, exceeding AF3 by over 10% of the total dataset. This is particularly valuable for peptide design, where high-quality structural models are essential. Considering the unique aspect of our noise initialization method, we further stratified the DockQ analysis based on peptide sequence length into three groups: 5-9, 10-15, and 16-30 residues (Fig. 3d and Extended Data Fig. 3). This analysis reveals that SiteAF3 shows a substantial improvement over AF3 for the shorter peptide lengths (5-9 and 10-15 residues), with a modest improvement for the 16-30 residue group. This observation aligns with the intuition behind our noise initialization strategy: shorter peptides, having a more confined spatial distribution, benefit more from a focused initialization around the binding pocket center, whereas longer peptides often have more extensive interaction surfaces with the protein, making a centralized initialization less optimal.

The case studies presented in Fig. 3E and Extended Data Fig. 3E and 3F further illustrate the superior performance of SiteAF3 over AF3 in three distinct ways. In the case of 8tmz, AF3 encountered issues during the prediction of the antibody structure, leading to a distorted pocket environment and hindering the accurate prediction of peptide binding. SiteAF3, however, demonstrated excellent performance. For 8p0q and 8ia5 [20], although AF3 predicted accurate protein structures, it completely misjudged the peptide binding sites, whereas SiteAF3 successfully ensured the peptide bound to the correct locations on the receptor proteins. Furthermore, in 8s6n [21], AF3 incorrectly folded a random coil region at the C-terminus of the full-length protein into a *β*-sheet, which occupied the original peptide binding pocket, directly leading to a failed structure prediction. Intriguingly, when the full-length sequence was not used, AF3 was able to successfully predict the complex structure. In contrast, SiteAF3 does not exhibit this sensitivity to potential mismatches between the sequence length and the crystal structure.

### 2.4 Results on protein and nucleic acids complexes

Aptamers represent another class of high-potential protein binders [22]. Deciphering the intricate interactions between nucleic acids and proteins is paramount for the rational design of effective aptamers. SiteAF3 has the capability to further enhance the prediction accuracy of protein-nucleic acid complexes. Consistent with the original AF3 paper, we primarily focused on the interactions between proteins and doublestranded DNA (dsDNA) as well as proteins and RNA. The protein-dsDNA dataset was compiled from the protein-DNA dataset available in FoldBench, while the protein-RNA dataset was identical to the one used in the original AF3 study (Fig. 4 and Extended Data Table 1). Our evaluation included the same four configuration modes of SiteAF3.

**Fig. 4:**
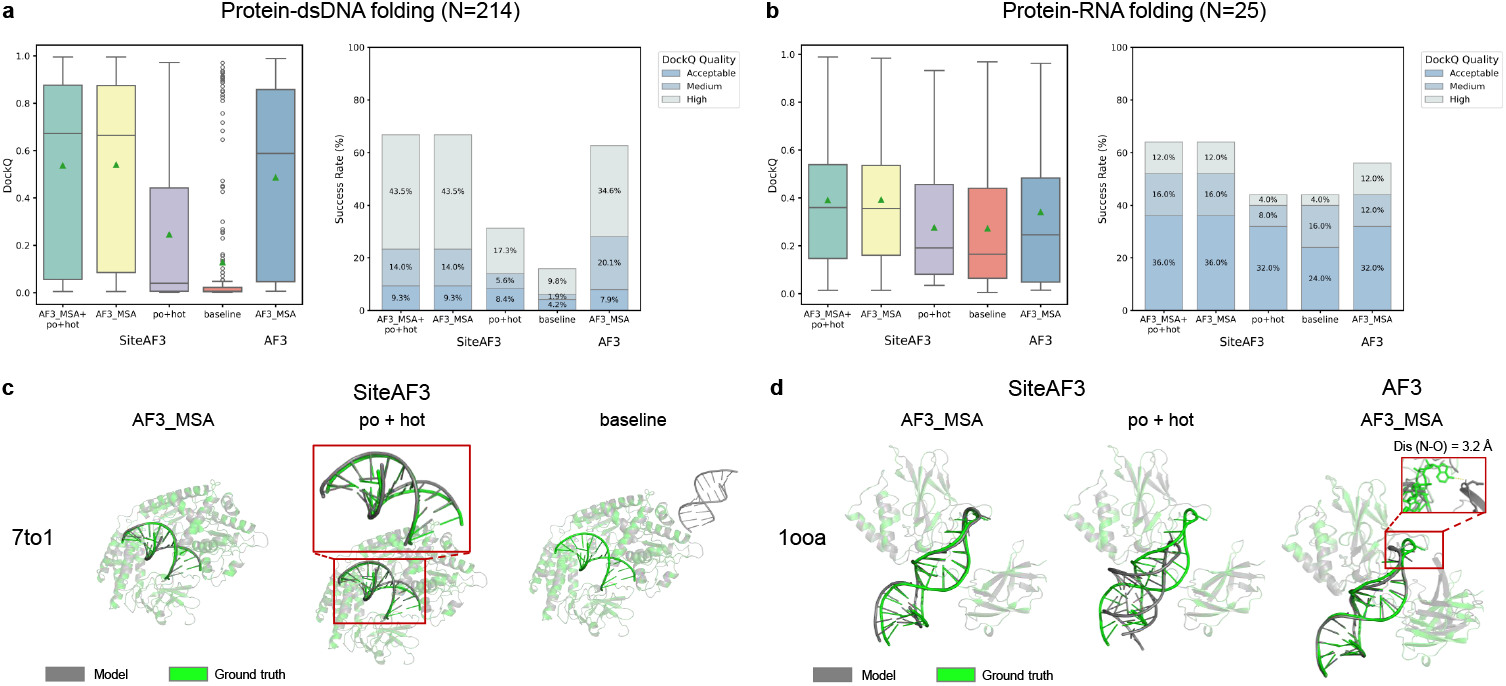
Results of protein and nucleic acids complexes. Each sample is assigned 3 random numbers, and each random number has 5 samples. The statistic analysis adopted the best results among these 15 structures. **a**, Docking performance of protein-dsDNA. **b**, Docking performance of protein-RNA. **c**, Case study: RIG-I bound to the end of p3SLR30 (PDB 7to1). All the three models were predicted by SiteAF3, with AF3 MSA succeeded (Mode 2), pocket and hotspot information (Mode 3) enabled dsDNA at the right site but with a base pair shift, and the baseline model (Mode 4) put dsDNA at wrong places. **d**, Case study: NF-kappaB(p50)2 complexed with a RNA aptamer (PDB 1ooa). SiteAF3 AF3 MSA (Mode 2) succeeded in docking, pocket and hotspot information (Mode 3) helped locating the RNA, while AF3 failed to generate right conformation of certain region in the protein, leading to an incorrect orientation of the base.

Fig. 4a illustrates that SiteAF3, when employing the AF3 MSA mode, demonstrates a significant improvement in the prediction quality of protein-dsDNA complexes. The average DockQ score increased by 10.2%, and the median DockQ score improved by 13.6%. Notably, the inclusion of pocket and hotspot residue information also substantially enhances the structural prediction accuracy compared to the baseline model. Fig. 4b shows a similar trend for protein-RNA complexes, where SiteAF3, again in the AF3 MSA mode, exhibits a marked enhancement in prediction quality. The average DockQ score increased by 14.7%, and the median DockQ score saw a substantial rise of 44.0%. Furthermore, the incorporation of pocket and hotspot residue information provides a noticeable boost to the structural prediction accuracy compared to the baseline. We further investigated the relationship between the ranking score and DockQ (Extended Data Fig. 4a). For protein-dsDNA complexes, the two SiteAF3 modes utilizing AF3 MSA exhibit a trend similar to that of AF3, but with a consistently higher average DockQ quality. However, for RNA complexes, due to the smaller sample size, discerning a definitive trend proved challenging. Extended Data Fig. 4b and 4c highlight SiteAF3’s improved ability to accurately capture the conformations of nucleic acid molecules and the intricate interactions at the protein-nucleic acid interface. It is important to note that nucleic acid molecules tend to be considerably larger than peptides, which may not be ideally suited for our current noise initialization approach. The primary gains in model performance can likely be attributed to the enhanced accuracy of protein structure prediction.

All three models featured in the case study of 7to1 [23] in Fig. 4c are based on SiteAF3. The baseline model incorrectly positioned the dsDNA molecule. The inclusion of pocket and hotspot information enabled the dsDNA to be placed at the correct binding site but with a shift in base pairs. Only the AF3 MSA mode, leveraging the complete template information through genetic searching, succeeded in accurately predicting the complex structure. Fig. 4d presents an interesting example where AF3 failed to generate the correct conformation of a specific region within the protein, leading to an altered orientation of a base in the DNA [24]. While pocket and hotspot information were sufficient to locate the binding site, they provided limited guidance for the precise structure of the nucleic acid. Only the utilization of the complete template information through genetic searching allowed for the accurate refinement of the nucleic acid structure.

### 2.5 The advantages of SiteAF3 in computing cost

SiteAF3 demonstrates advantages over AF3 in terms of computing cost in two key aspects. The first pertains to GPU memory consumption (Extended Data Table 2). During our training, using an RTX 6000 Ada graphics card with 48 GB of memory, we observed that AF3 is prone to out-of-memory errors for structures with a total token count exceeding approximately 3500. In contrast, when using SiteAF3, the implementation of a mask that selectively cuts the two-pair embedding matrices during matrix multiplication to update only the ligand coordinates significantly reduces the memory footprint of this step. This allows for the computation of larger bio-systems. The second advantage arises when AF3 MSA is not used, and the prediction relies solely on pocket and hotspot information. While this configuration might lead to a reduction in accuracy in some cases, it avoids the computationally intensive genetic searching process, resulting in a substantial speedup in the overall runtime. Compared to the AF3 model without AF3 MSA, SiteAF3 in this mode offers acceptable accuracy. Thus, SiteAF3 in this mode is potentially valuable for high-throughput screening applications.

## 3 Conclusions

SiteAF3, a novel method for accurate site-specific folding based on conditional diffusion within the AF3 framework, demonstrates significant improvements in predicting the structures of diverse biomolecular complexes, especially for allosteric sites, flexible structures and orphan proteins with known structures. In the realm of protein-small molecule interactions, evaluations on the FoldBench and PoseBustersV2 datasets reveal that SiteAF3 consistently outperforms AF3, particularly when incorporating binding pocket and hotspot residue information. Notably, SiteAF3 achieves higher accuracy for less confident predictions and shows substantial reductions in Ligand RMSD. For protein-peptide complexes, tested on the PepPCBench dataset, SiteAF3 exhibits significantly higher DockQ scores and an enhanced ability to model both interface interactions and peptide conformations, especially for shorter peptides. Furthermore, SiteAF3 demonstrates superior performance in predicting protein-nucleic acid (dsDNA and RNA) complexes, as evidenced by substantial increases in average and median DockQ scores on dedicated datasets. This improvement is particularly pronounced when leveraging the MSA information from AF3. Beyond accuracy enhancements, SiteAF3 also presents computational advantages, including a reduced GPU memory footprint for large systems and a faster runtime when utilizing only pocket and hotspot information, offering a potential avenue for high-throughput screening.

Despite the promising results, SiteAF3 has some limitations. Firstly, the current noise initialization strategy for the ligand relies on a simplified approach of placing initial coordinates around the pocket center. This might not be optimal for all binding scenarios, particularly for ligands with more extended or complex binding interfaces. Future work could explore making the noise initialization learnable or adaptive based on the shape of the binding interface to improve the sampling efficiency. Secondly, SiteAF3 in its current form lacks a specific module for handling covalent docking. Expanding the functionality to accurately model covalent interactions would significantly broaden the applicability of SiteAF3 in drug discovery and the study of enzyme mechanisms.

## 4 Methods

### 4.1 Model Inference

The general workflow for SiteAF3 input generation is as follows: First, prepare a structure file in the .pdb format. Subsequently, preprocess the structure file using PDBFixer [25] or pdb4amber [26] to repair any missing atoms and residues. Then, generate the corresponding hotspot and pocket structure files using the *generate hotspot*.*py* and *generate pocket*.*py* scripts. Alternatively, the same operations can be performed using PyMOL [27]. The pocket file is essential for the noise initialization step. Finally, generate the corresponding JSON input files based on the receptor and ligand information.

For the tests presented in this study, we first preprocessed the files for all datasets to obtain the ground truth structures. We identified the location of the target ligand and then selected all receptor chains within a 10 Å radius of the ligand, packaging them together into a complex structure. This approach effectively controlled the total number of tokens in the input and minimized potential impacts on the binding site. When generating the JSON input files for both SiteAF3 and AF3, we did not directly extract sequence information from the ground truth structures. Instead, we retrieved the sequence information based on the respective chain IDs from FASTA files or from the chemical component dictionary (CCD) downloaded from the RCSB PDB website.

After input generation, predictions for protein-small molecule datasets were performed using a 5×5 sampling strategy (5 seeds × 5 samples). Other datasets utilized a 3×5 sampling strategy (3 seeds × 5 samples). Unless otherwise specified, the hotspot residue truncation radius is 7 Å, and the pocket residue truncation radius is 10 Å near the hotspot residues. Both SiteAF3 and AF3 employed the genetic searching tool provided by the official AF3 release. The majority of inference runs were conducted on 48 GB NVIDIA RTX 6000 Ada GPUs. For specific cases where AF3 encountered outof-memory errors, inference was performed using NVIDIA A100 GPUs with 80 GB of memory.

### 4.2 Evaluation and Metrics

The primary software packages utilized for analysis were PyMOL 3.1.0 and DockQ v2. For protein-small molecule complexes, Ligand RMSD (LRMSD) was primarily measured using PyMOL. Given that the atom numbering within the small molecule can change in the conformations output by SiteAF3 and AF3, we implemented a specific processing step. Firstly, we performed a structural superposition by aligning the binding site of the receptor. Subsequently, we employed a greedy algorithm based on atom types to match the ligand atoms and calculate the RMSD. Consistent with the AF3 paper, the ligand docking success rate is defined as LRMSD *<* 2 Å.

For protein-peptide complexes, we used DockQ v2 to assess the quality of the interaction between the designated receptor chain and the peptide. It is important to note that when multiple receptor chains were present in the same structure, we adopted a similar approach to DockQ v1, whereby we calculated the DockQ score for the peptide against each individual receptor chain and then reported the average score.

For protein-nucleic acid complexes, we again utilized DockQ v2 for quality assessment. However, in contrast to peptide complexes, the authors of the FoldBench dataset separately reported the score for each receptor chain against each nucleic acid chain. We followed the same convention in our analysis. To provide a clearer assessment of docking results, DockQ scores can be categorized into four bins based on their values:

- Incorrect: DockQ *<* 0.23
- Acceptable: 0.23 *<*= DockQ *<* 0.49
- Medium: 0.49 *<*= DockQ *<* 0.80
- High: DockQ *>* 0.80

## Code availability

All the source code is available on https://github.com/HaCTang/SiteAF3 or https://github.com/ClickFF/SiteAF3.

## Acknowledgements

This work was supported by funds from the National Institutes of Health (R01GM147673, R01GM149705) and the National Science Foundation (1955260). The authors would like to thank the computing resources provided by the Center for Research Computing (facility RRID: SCR 022735) at the University of Pittsburgh (NSF award number OAC-2117681), and the Pittsburgh Supercomputer Center (grant number BIO210185).

The following algorithm illustrates the realization of core module in SiteAF3. Any other functions applied in the algorithm inherit from AF3.

### Algorithm 1

Conditional Pocket Diffusion Sampling

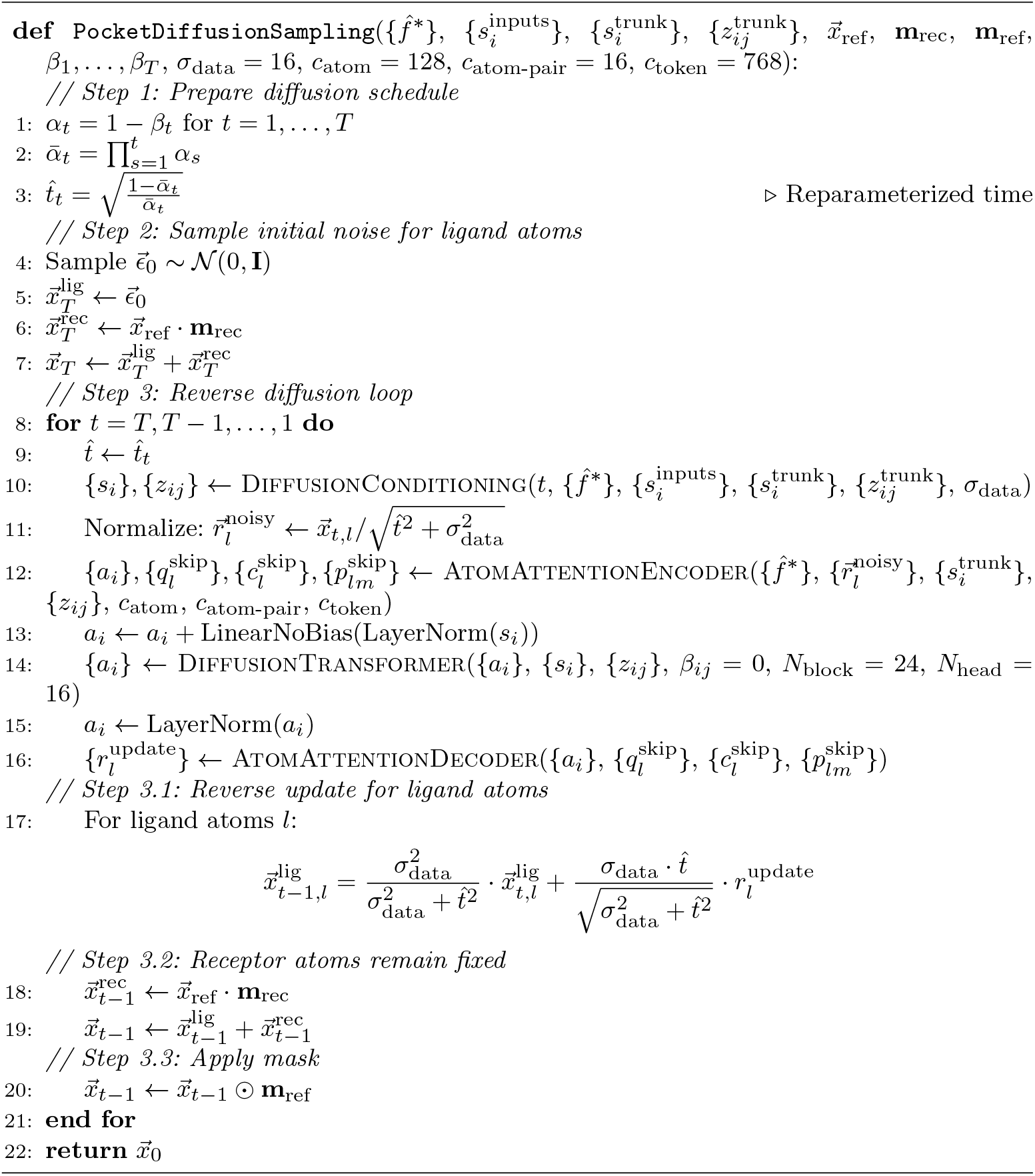

---

**Extended Data Fig 1.**
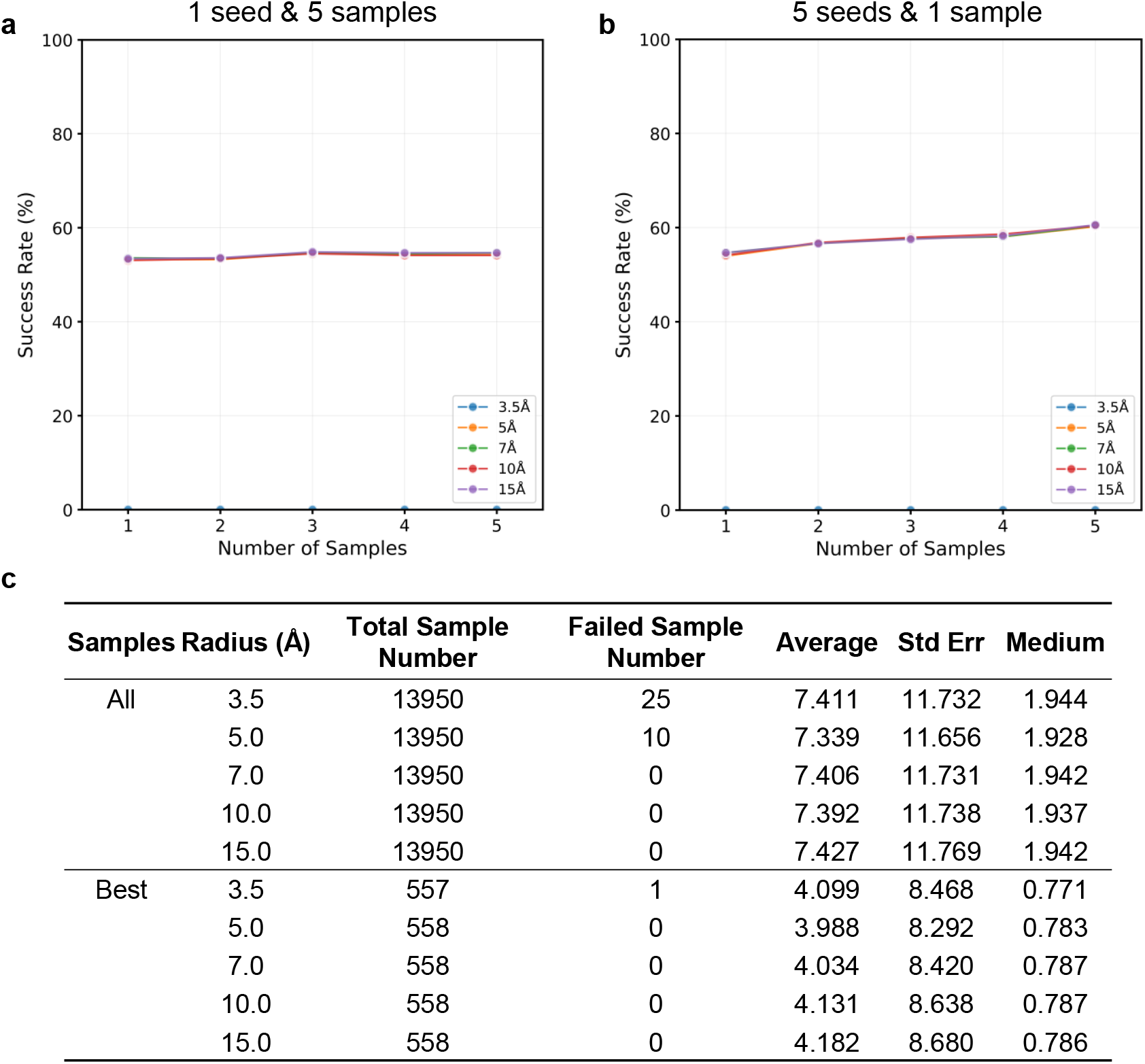
Supplementary influence of noise initialization radius on the structure prediction of protein-ligand complexes. All results are based on 5 radius, 3.5 Å, 5.0 Å, 7.0 Å, 10.0 Å, 15.0 Å. **a, b**, The individual impact of the number of seeds and samples on performance. **c**, Statistical values of sample radius evaluation.

**Extended Data Fig 2.**
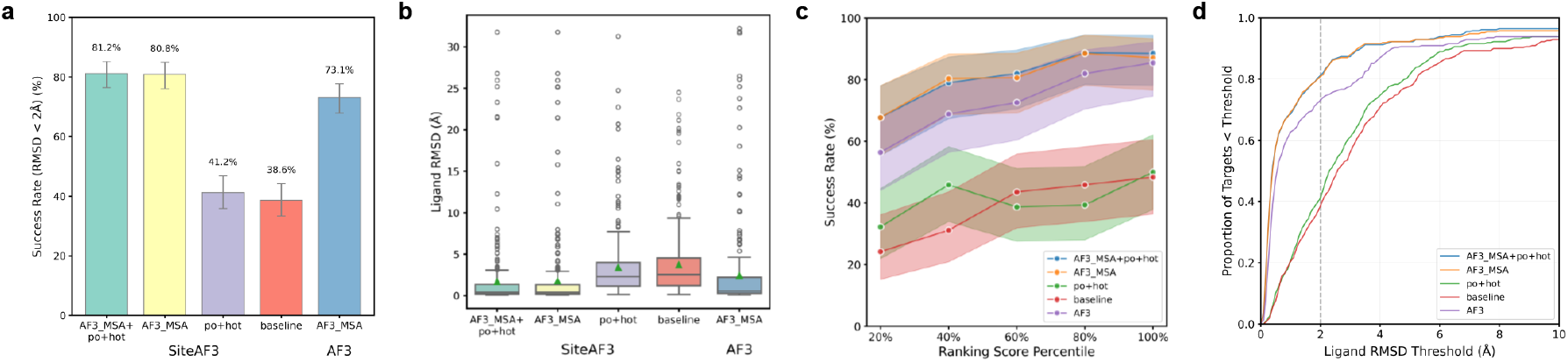
Results of protein and small molecule complexes on PoseBustersV2 datasets. Each sample is assigned 5 random numbers, and each random number has 5 samples. The statistic analysis adopted the best results among these 25 structures. **a**, Success rate across different subsets, where the success rate is defined as the ratio of LRMSD less than 2 Å. **b**, Comparison of ligand RMSD between SiteAF3 and AF3. **c**, The relationship between success rate and confidence-related scores of SiteAF3 and AF3. The scores were segmented by number quantiles, with each bin evenly dividing our test samples. **d**, Cumulative density of ligand RMSD, varying from 0 to 10 Å.

**Extended Data Fig 3.**
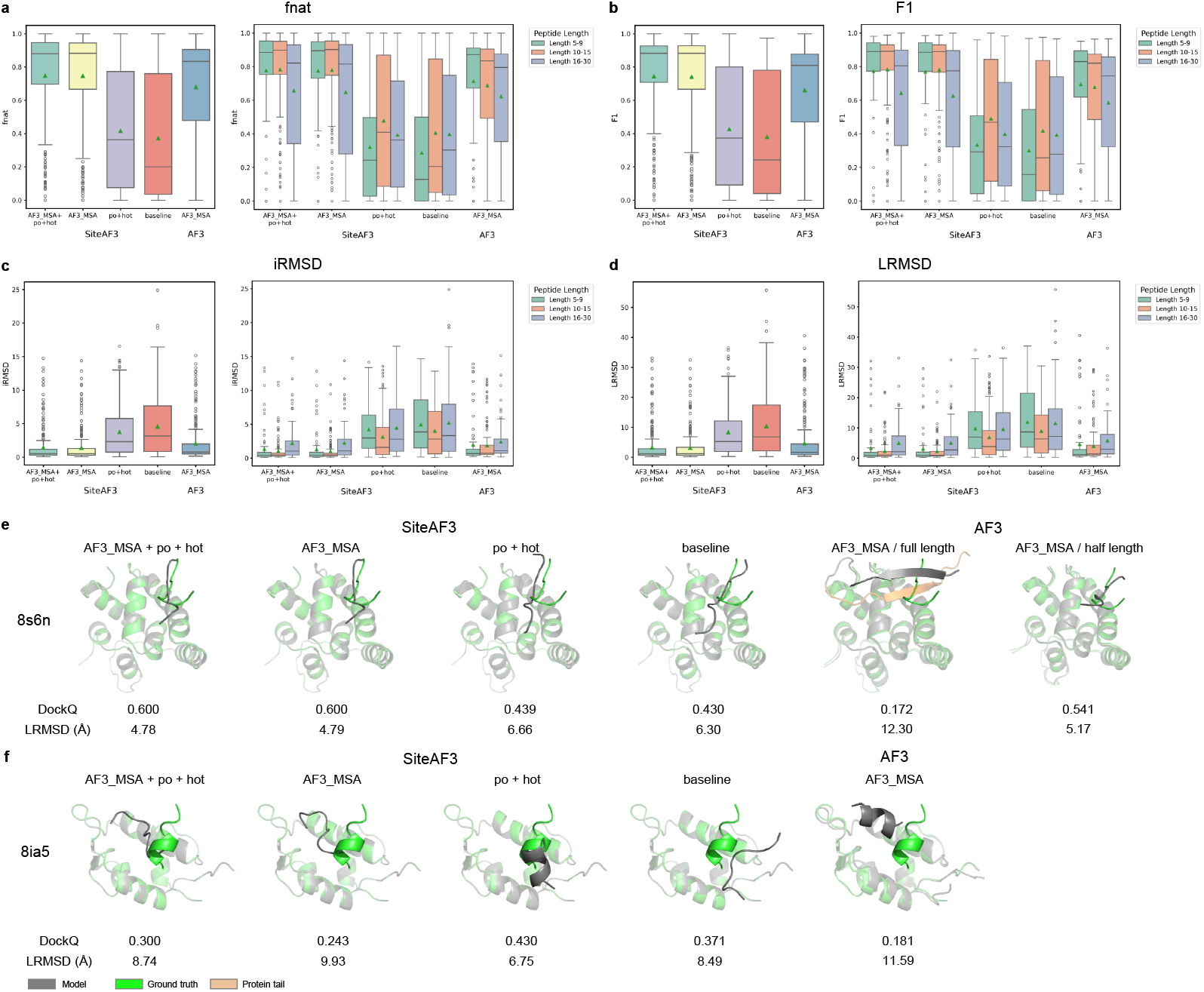
Supplementary results of protein and peptide complexes on PepPCBench datasets. **a**, Fraction of native contact (fnat) analysis on all (left) and length-devided (right) datasets between SiteAF3 and AF3. **b**, Harmonic mean of the precision and recall in predicted interfacial contact (F1) analysis on all (left) and length-devided (right) datasets between SiteAF3 and AF3. **c**, iRMSD analysis on all (left) and length-devided (right) datasets between SiteAF3 and AF3. **d**, LRMSD analysis on all (left) and length-devided (right) datasets between SiteAF3 and AF3. **e**, Case study: MLLE3 domain of Rrm4 in complex with PAM2L1 of Upa1 (PDB 8s6n). All 4 modes in SiteAF3 succeeded in docking the peptide. AF3 failed to generate right conformation of peptide when using full length sequnce of proteins due to wrong folding of terminal random coil. **f**, Case study: Small peptide binds to N-terminal domain of MdmX. When the receptor protein successfully folds, the docking result of AF3 is worse than that of SiteAF3.

**Extended Data Fig 4.**
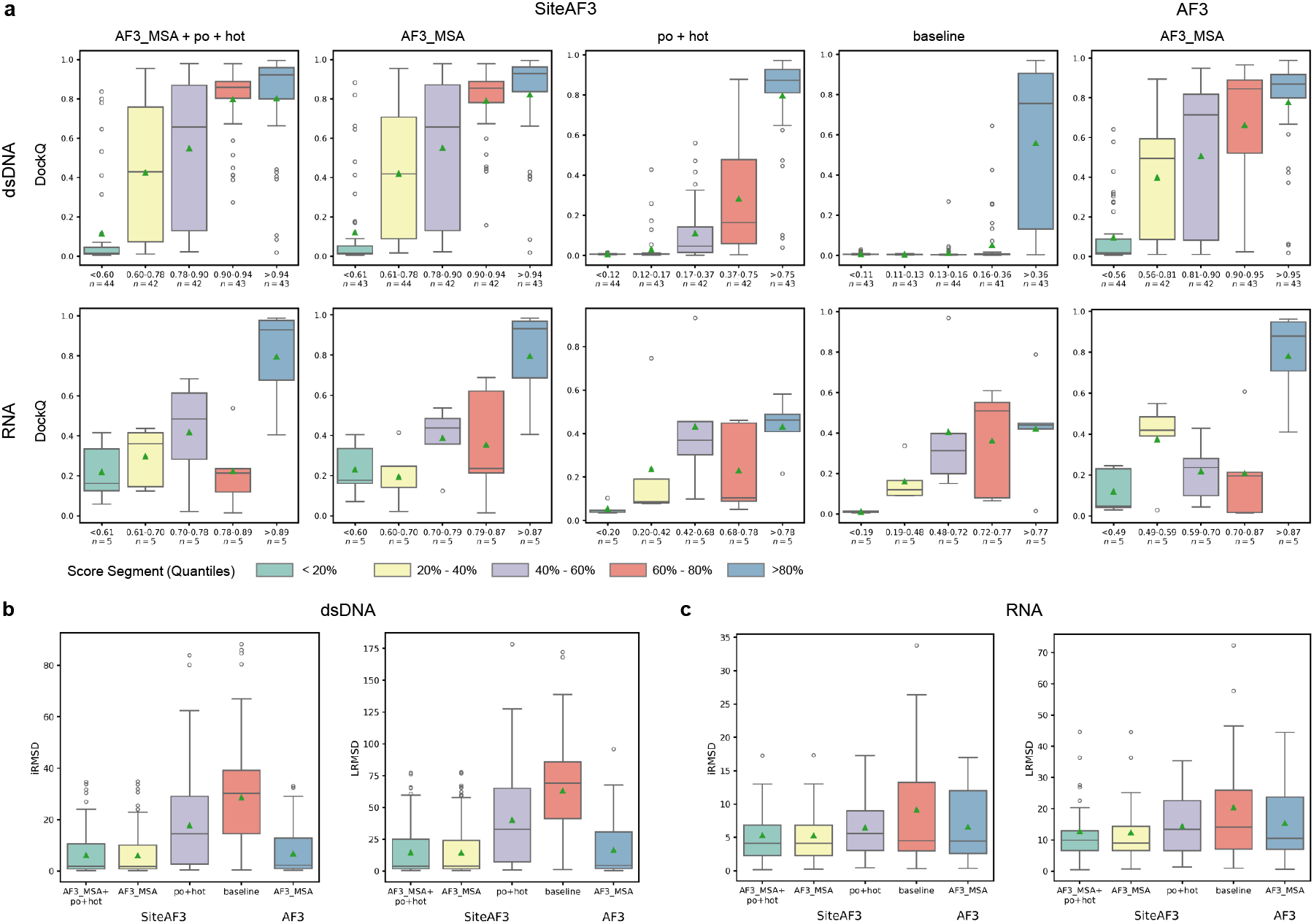
Supplementary results of protein and nucleic acid complexes. **a**, The relationship between DockQ and confidence-related scores. The scores were segmented by number quantiles, with each bin evenly dividing our test samples. Up: protein-dsDNA. Down: protein-RNA. **b**, iRMSD and LRMSD analysis of protein-dsDNA complexes. **c**, iRMSD and LRMSD analysis of protein-RNA complexes.

**Extended Data Table 2:** Case analysis of GPU memory consumption.

| Input Matrix | Shape | Note |
| --- | --- | --- |
| A | (3500, 3500, 128) |  |
| B | (3500, 3500, 128) |  |
| mask | (3500,) | Suppose the first 2000 bits are 0 and the last 1500 bits are 1 |
| A' | (1500, 3500, 128) | A tailored by mask |
| B' | (1500, 3500, 128) | B tailored by mask |
| Calculation | Memory (float16) |  |
| A*B=C | ~ 9.4 GB |  |
| A'*B'=C' | ~ 4.1 GB |  |

## Notes

### Competing Interest Statement

The authors have declared no competing interest.

## References

[1] Jumper, J., Evans, R., Pritzel, A., Green, T., Figurnov, M., Ronneberger, O., Tunyasuvunakool, K., Bates, R., Žídek, A., Potapenko, A., Bridgland, A., Meyer, C., Kohl, S.A.A., Ballard, A.J., Cowie, A., Romera-Paredes, B., Nikolov, S., Jain, R., Adler, J., Back, T., Petersen, S., Reiman, D., Clancy, E., Zielinski, M., Steinegger, M., Pacholska, M., Berghammer, T., Bodenstein, S., Silver, D., Vinyals, O., Senior, A.W., Kavukcuoglu, K., Kohli, P., Hassabis, D.: Highly accurate protein structure prediction with alphafold. Nature 596(7873), 583–589 (2021) 10.1038/s41586-021-03819-2. [Online; accessed 2025-07-04]

[2] Jumper, J., Evans, R., Pritzel, A., Green, T., Figurnov, M., Ronneberger, O., Tunyasuvunakool, K., Bates, R., Žídek, A., Potapenko, A., Bridgland, A., Meyer, C., Kohl, S.A.A., Ballard, A.J., Cowie, A., Romera-Paredes, B., Nikolov, S., Jain, R., Adler, J., Back, T., Petersen, S., Reiman, D., Clancy, E., Zielinski, M., Steinegger, M., Pacholska, M., Berghammer, T., Bodenstein, S., Silver, D., Vinyals, O., Senior, A.W., Kavukcuoglu, K., Kohli, P., Hassabis, D.: Highly accurate protein structure prediction with alphafold. Nature 596(7873), 583–589 (2021) 10.1038/s41586-021-03819-2. [Online; accessed 2025-07-04]

[3] Rosignoli, S., Pacelli, M., Manganiello, F., Paiardini, A.: An outlook on structural biology after <scp>a</scp>lpha<scp>f</scp>old: tools, limits and perspectives. FEBS Open Bio 15(2), 202–222 (2024) 10.1002/2211-5463.13902. [Online; accessed 2025-07-04]

[4] Liu, J., Neupane, P., Cheng, J.: Boosting AlphaFold Protein Tertiary Structure Prediction through MSA Engineering and Extensive Model Sampling and Ranking in CASP16. Cold Spring Harbor Laboratory. [Online; accessed 2025-07-04] (2025). 10.1101/2025.06.06.658338

[5] Michaud, J.M., Madani, A., Fraser, J.S.: A language model beats alphafold2 on orphans. Nature Biotechnology 40(11), 1576–1577 (2022) 10.1038/s41587-022-01466-0. [Online; accessed 2025-07-04]

[6] Nittinger, E., Yoluk,, Tibo, A., Olanders, G., Tyrchan, C.: Co-folding, the future of docking – prediction of allosteric and orthosteric ligands. Artificial Intelligence in the Life Sciences, 100136 (2025) 10.1016/j.ailsci.2025.100136. [Online; accessed 2025-07-04]

[7] La, H., Gupta, A., Morehead, A., Cheng, J., Zhang, M.: MegaFold: System-Level Optimizations for Accelerating Protein Structure Prediction Models (2025). https://arxiv.org/abs/2506.20686

[8] Wang, F., Wang, Y., Feng, L., Zhang, C., Lai, L.: Target-specific De Novo peptide binder design with diffpepbuilder. Journal of Chemical Information and Modeling 64(24), 9135–9149 (2024) 10.1021/acs.jcim.4c00975. [Online; accessed 2025-07-04]

[9] Xu, S., Feng, Q., Qiao, L., Wu, H., Shen, T., Cheng, Y., Zheng, S., Sun, S.: FoldBench: An All-atom Benchmark for Biomolecular Structure Prediction. Cold Spring Harbor Laboratory. [Online; accessed 2025-07-04] (2025). 10.1101/2025.05.22.655600

[10] Buttenschoen, M., Morris, G.M., Deane, C.M.: Posebusters: Ai-based docking methods fail to generate physically valid poses or generalise to novel sequences. Chemical Science 15(9), 3130–3139 (2024) 10.1039/d3sc04185a. [Online; accessed 2025-07-04]

[11] Brear, P., De Fusco, C., Atkinson, E.L., Iegre, J., Francis-Newton, N.J., Venkitaraman, A.R., Hyvönen, M., Spring, D.R.: A fragment-based approach leading to the discovery of inhibitors of ck2 with a novel mechanism of action. RSC Medicinal Chemistry 13(11), 1420–1426 (2022) 10.1039/d2md00161f. [Online; accessed 2025-07-04]

[12] Steinebach, C., Bricelj, A., Murgai, A., Sosič, I., Bischof, L., Ng, Y.L.D., Heim, C., Maiwald, S., Proj, M., Voget, R., Feller, F., Košmrlj, J., Sapozhnikova, V., Schmidt, A., Zuleeg, M.R., Lemnitzer, P., Mertins, P., Hansen, F.K., Gütschow, M., Krönke, J., Hartmann, M.D.: Leveraging ligand affinity and properties: Discovery of novel benzamide-type cereblon binders for the design of protacs. Journal of Medicinal Chemistry 66(21), 14513–14543 (2023) 10.1021/acs.jmedchem.3c00851. [Online; accessed 2025-07-04]

[13] Yin, S., Mi, X., Shukla, D.: Leveraging machine learning models for peptide–protein interaction prediction. RSC Chemical Biology 5(5), 401–417 (2024) 10.1039/d3cb00208j. [Online; accessed 2025-07-04]

[14] Evans, R., O’Neill, M., Pritzel, A., Antropova, N., Senior, A., Green, T., Žídek, A., Bates, R., Blackwell, S., Yim, J., Ronneberger, O., Bodenstein, S., Zielinski, M., Bridgland, A., Potapenko, A., Cowie, A., Tunyasuvunakool, K., Jain, R., Clancy, E., Kohli, P., Jumper, J., Hassabis, D.: Protein complex prediction with AlphaFold-Multimer. Cold Spring Harbor Laboratory. [Online; accessed 2025-07-04] (2021). 10.1101/2021.10.04.463034

[15] Zhai, S., Zhao, H., Wang, J., Lin, S., Liu, T., Jiang, D., Liu, H., Kang, Y., Yao, X., Hou, T.: PepPCBench is a Comprehensive Benchmark for Protein-Peptide Complex Structure Prediction with AlphaFold3. Cold Spring Harbor Laboratory. [Online; accessed 2025-07-04] (2025). 10.1101/2025.04.08.647699

[16] Mirabello, C., Wallner, B.: Dockq v2: improved automatic quality measure for protein multimers, nucleic acids, and small molecules. Bioinformatics 40(10) (2024) 10.1093/bioinformatics/btae586. [Online; accessed 2025-07-04]

[17] Jandova, Z., Vargiu, A.V., Bonvin, A.M.J.J.: Native or non-native protein–protein docking models? molecular dynamics to the rescue. Journal of Chemical Theory and Computation 17(9), 5944–5954 (2021) 10.1021/acs.jctc.1c00336. [Online; accessed 2025-07-04]

[18] Shirali, A., Stebliankin, V., Karki, U., Shi, J., Chapagain, P., Narasimhan, G.: A comprehensive survey of scoring functions for protein docking models. BMC bioinformatics 26(1), 25 (2025) 10.1186/s12859-024-05991-4. [Online; accessed 2025-07-04]

[19] Piniello, B., Macías-León, J., Miyazaki, S., García-García, A., Compañón, I., Ghirardello, M., Taleb, V., Veloz, B., Corzana, F., Miyagawa, A., Rovira, C., Hurtado-Guerrero, R.: Molecular basis for bacterial n-glycosylation by a soluble hmw1c-like n-glycosyltransferase. Nature Communications 14(1) (2023) 10.1038/s41467-023-41238-1. [Online; accessed 2025-07-04]

[20] Cheng, X., Chen, R., Zhou, T., Zhang, B., Li, Z., Gao, M., Huang, Y., Liu, H., Su, Z.: Leveraging the multivalent p53 peptide-mdmx interaction to guide the improvement of small molecule inhibitors. Nature Communications 13(1) (2022) 10.1038/s41467-022-28721-x. [Online; accessed 2025-07-04]

[21] Devan, S.-K., Shanmugasundaram, S., Müntjes, K., Postma, J., Smits, S.H.J., Altegoer, F., Feldbrügge, M.: Deciphering the rna-binding protein network during endosomal mrna transport. Proceedings of the National Academy of Sciences 121(46) (2024) 10.1073/pnas.2404091121. [Online; accessed 2025-07-04]

[22] Zhou, J., Rossi, J.: Aptamers as targeted therapeutics: current potential and challenges. Nature reviews. Drug discovery 16(3), 181–202 (2017) 10.1038/nrd.2016.199. [Online; accessed 2025-07-04]

[23] Wang, W., Pyle, A.M.: The rig-i receptor adopts two different conformations for distinguishing host from viral rna ligands. Molecular Cell 82(21), 4131–41446 (2022) 10.1016/j.molcel.2022.09.029. [Online; accessed 2025-07-04]

[24] Huang, D.-B., Vu, D., Cassiday, L.A., Zimmerman, J.M., Maher, L.J., Ghosh, G.: Crystal structure of nf-b (p50) 2 complexed to a high-affinity rna aptamer. Proceedings of the National Academy of Sciences 100(16), 9268–9273 (2003) 10.1073/pnas.1632011100. [Online; accessed 2025-07-04]

[25] Eastman, P., Swails, J., Chodera, J.D., McGibbon, R.T., Zhao, Y., Beauchamp, K.A., Wang, L.-P., Simmonett, A.C., Harrigan, M.P., Stern, C.D., Wiewiora, R.P., Brooks, B.R., Pande, V.S.: Openmm 7: Rapid development of high performance algorithms for molecular dynamics. PLoS computational biology 13(7), 1005659 (2017) 10.1371/journal.pcbi.1005659. [Online; accessed 2025-07-04]

[26] Case, D.A., Aktulga, H.M., Belfon, K., Cerutti, D.S., Cisneros, G.A., Cruzeiro, V.W.D., Forouzesh, N., Giese, T.J., Götz, A.W., Gohlke, H., Izadi, S., Kasavajhala, K., Kaymak, M.C., King, E., Kurtzman, T., Lee, T.-S., Li, P., Liu, J., Luchko, T., Luo, R., Manathunga, M., Machado, M.R., Nguyen, H.M., O’Hearn, K.A., Onufriev, A.V., Pan, F., Pantano, S., Qi, R., Rahnamoun, A., Risheh, A., Schott-Verdugo, S., Shajan, A., Swails, J., Wang, J., Wei, H., Wu, X., Wu, Y., Zhang, S., Zhao, S., Zhu, Q., Cheatham, T.E., Roe, D.R., Roitberg, A., Simmerling, C., York, D.M., Nagan, M.C., Merz, K.M.: Ambertools. Journal of Chemical Information and Modeling 63(20), 6183–6191 (2023) 10.1021/acs.jcim.3c01153. [Online; accessed 2025-07-04]

[27] Schrödinger, L., DeLano, W.: PyMOL. http://www.pymol.org/pymol

